# The F pilus serves as a conduit for the DNA during conjugation between physically distant bacteria

**DOI:** 10.1101/2023.06.21.545889

**Authors:** Kelly Goldlust, Adrien Ducret, Manuel Halte, Annick Dedieu-Berne, Marc Erhardt, Christian Lesterlin

**Affiliations:** Molecular Microbiology and Structural Biochemistry (MMSB), Université Lyon 1, CNRS, Inserm, UMR5086, 69007, Lyon, France; Institute for Biology/Molecular Microbiology, Humboldt-Universität zu Berlin, 10115, Berlin, Germany; Max Planck Unit for the Science of Pathogens, 10117, Berlin, Germany

**Keywords:** Horizontal gene transfer, Bacterial conjugation, F Pilus

## Abstract

Horizontal transfer of F-like plasmids by bacterial conjugation is responsible for disseminating antibiotic resistance and virulence determinants among pathogenic Enterobacteriaceae species, a growing health concern worldwide. Central to this process is the conjugative F pilus, a long extracellular filamentous polymer that extends from the surface of plasmid donor cells, allowing it to probe the environment and make contact with the recipient cell. It is well established the F pilus can retract to bring mating pair cells in tight contact before DNA transfer. However, whether DNA transfer can occur through the extended pilus has been a subject of active debate. In this study, we use live-cell microscopy to show that the F pilus can indeed serve as a conduit for the DNA during transfer between physically distant cells. Our findings enable us to propose a new model for conjugation that revises our understanding of the DNA transport mechanism and the dissemination of drug resistance in complex bacterial communities.

**One-Sentence Summary:** Plasmid DNA passes through the F pilus during conjugational transfer between physically distant bacteria.

## Introduction

Bacterial conjugation is a contact-dependent horizontal gene transfer mechanism that plays a crucial role in the evolution of bacterial genomes and the dissemination of metabolic properties, such as resistance to antibiotics, heavy metals and virulence (Lederberg and Tatum, 1946; Lawrence, 1999; Thomas and Nielsen, 2005; Barlow, 2009; Virolle *et al*., 2020). Paradigmatic examples are members of the conjugative F plasmid family, which are highly transmissible and prevalent among clinical isolates of enterobacterial pathogens, including *Escherichia coli*, *Klebsiella pneumoniae, Salmonella enterica* and *Shigella flexneri* (Garcillán-Barcia *et al*., 2011; Koraimann, 2018; Stephens *et al*., 2020). Conjugational transfer of F-like plasmids requires a Type-IV secretion system (T4SS) that transports the single-stranded DNA (ssDNA) plasmid through the cell envelop and produces the conjugative pilus, which is also strictly essential for conjugation. The structure and function of F-like conjugative pili have been extensively studied, particularly those encoded by the F and the pED208 plasmids. The F pilus is an assembly of thousands of TraA pilin (VirB2) monomers polymerized into a long, flexible and tubular helical filament 87 Å in width and an average ∼1-2 µm in length, which extends from the surface of the donor cells (Date *et al*., 1977; Bradley, 1984; Paranchych and Frost, 1988; Costa *et al*., 2016; Kishida *et al*., 2022). Notably, the pilus is a highly dynamic structure that undergoes cycles of extension and retraction (Daehnel *et al*., 2005; Clarke *et al*., 2008; Ilangovan *et al*., 2015; Goldlust *et al*., 2022). The function of retraction and its requirement for conjugation have been actively debated. Using live-cell imaging of the F pilus labelled with fluorescent R17 bacteriophages, Clarke et al. (Clarke *et al*., 2008) showed that the F pilus randomly extends and retracts to scan the host cell’s surroundings to makes contact with a recipient cell. The subsequent retraction of the pilus establishes physical contact between the donor and recipient cells, thereby allowing the stabilization of the mating pair through interaction between the donor’s outer membrane protein TraN, and the recipient’s outer membrane protein OmpA (Achtman, 1975; Klimke and Frost, 1998; Klimke *et al*., 2005; Low *et al*., 2022). However, whether the extended pilus can serve as a conduit for the DNA during conjugation between cells that are physically distant remains an open question. This possibility was first proposed by Charles C. Brinton sixty years ago (Brinton et al., 1964; Brinton, 1965) and has since been a subject of extensive debate.

Little experimental evidence exist that support the possibility DNA transport through the extended pilus. Harrington and Rogerson (Harrington and Rogerson, 1990) reported conjugation between donor and recipient cells cultures separate by a filter 6 µm thick with pores 0.01 to 0.1 µm in diameter. Using live-cell microscopy and an elegant reporter system for DNA transfer, work by Babić et al. (Babic *et al*., 2008) provides evidence for DNA acquisition by recipients that are not in direct contact with a donor cell. However, their experimental setup could not exclude the possibility that conjugation had occurred in the conjugation mix before imaging and the extended pilus between donor and recipient cells was not visualized. The lack of direct evidence for distant transfer favoured the currently prevailling model that conjugation strictly requires tight contact between donor and recipient cells and the possibility that the DNA passes through the pilus lumen remains speculative In this study, we developped real-time fluorescence microscopy approach to perform the direct visualisation of the fluorescently labelled F pilus (Goldlust *et al*., 2022) and the transferred DNA during conjugation in live-cells (Nolivos *et al*., 2019; Couturier *et al*., 2023). Our results unequivocally demonstrate that the F pilus has a dual function in establishing cell-to-cell contact and serving as a conduit for DNA during transfer between physically distant cells.

## Results

### Cell piliation and F pilus morphology

The labelling of the F pilus was performed in the wild type (*wt*) *Escherichia coli* K12 MG1655 strain carrying the full length F plasmid and an expression plasmid (p *traA*^S54C^) coding for a mutant TraA^S54C^ pilin where the surface-exposed serine 54 was substituted by a cysteine for bioconjugation with Alexa Fluor-Maleimide (AF-Mal) (Fig. 1A) (Goldlust *et al*., 2022). This *wt E. coli* / F / p *traA*^S54C^ partial diploid strain produces both the *wt* TraA and the mutant TraA^S54C^ pilins and exhibits conjugation efficiency comparable to the *wt* F+ donor, both in the presence or the absence of maleimide labelling (Fig. S1A). Live-cell fluorescence microscopy imaging of the *wt* F / p *traA*^S54C^ strain after incubation with AF-Mal^488^ revealed efficient fluorescent labelling of the F pilus at the surface of the cells (Fig. 1B). To enable automatic pili detection, tracking and quantitative analysis, we developed a custom function for the MicrobeJ pluging (Ducret *et al*., 2016) that detects the pili anchoring points on the cell periphery and the shape of the extracellular pilus filament (Fig. 1B; Fig. S1B; Movie S1; Supplementary information).

**Figure 1.**
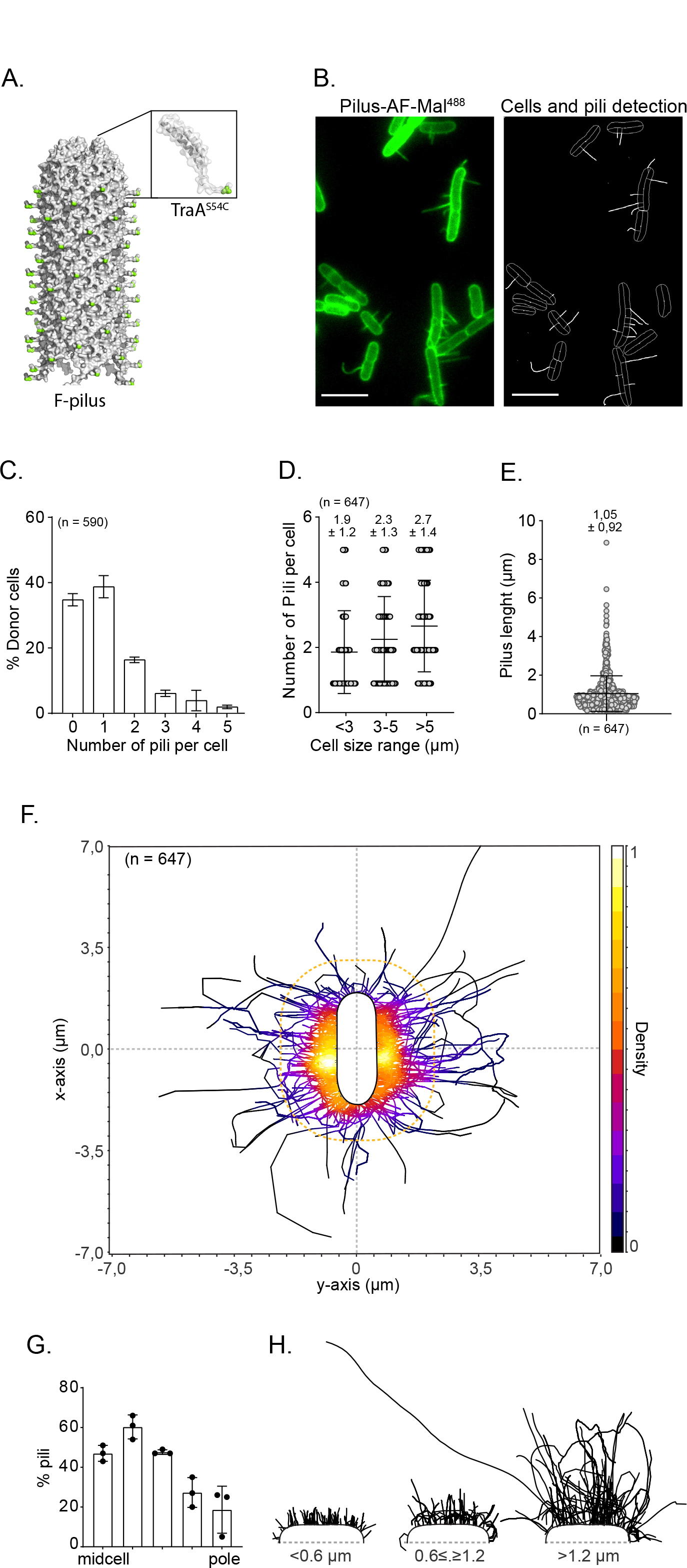
Characterisation of donor cells piliation (**A**) Surface representation map of the cryo-EM structure of the conjugative F-pilus (adapted form (Costa *et al*., 2016) to which the surface-exposed serine 54 (serine 3 of the mature pilin) replaced by a cysteine in the TraA^S54C^ mutant was added and displayed in green. The inset shows the structure of an individual TraA pilin to which the N-terminal carrying the serine 54 was added. (**B**) Microscopy image showing pili labelled with AF488-maleimide at the surface of F donor cells (left) and the corresponding detection of the cells outlines and fluorescent pili using MicrobeJ custom plugin (right). (**C**) Histogram of the number of pili per cells. The Mean and SD were calculated from the indicated number (n) of cells from three independent experiments. (**D**) Dot-plot of the number of pili per cell sorted by cell size range. Each open circle represents one cell. The Mean and SD were calculated from the indicated number of pili from three independent experiments. (**E**) Dot-plot of of pili length distribution. Mean and SD were calculated from the indicated number of Pili from three independent snapshot microscopy experiment (**F**) 2D density map of pili localisation around the cells normalised by the cell length. The number of pili (n) from three biological replicates is indicated. Density scale on the left. (**G**) Histogram of pili distribution along the cell perimeter from midcell to the cell tip. (**H**) Representation of the curvature of pili categorized by size range.

Analysis of snapshot microscopy images reveals that the number of pili per cell ranges from 0 to 5 (n = 590), with the majority of piliated cells harboring one (46,70 %) or two pili (19,67 %) (Fig. 1C). The number of pili per cell slightly increases from 1.9 ± 1.2 in to 2.7 ± 1.7 (n = 647) as the cell progresses through the cell cycle from newborn to pre-dividing cells (Fig. 1D). Analysis of pili length distribution shows that most pili are smaller than 2 µm in length and rarely exceed 4 µm, with an average length of 1.05 ± 0.92 µm (n = 647) (Fig. 1E). There is no significant correlation between the pili length and the number of pili per cells or the cell size (Fig. S1C-1D), indicating that the cells produce pili of similar length throughout the cell cycle, regardless of the number of pili per cell. This description of cell piliation is in good agreement with previous works using either maleimide bioconjugation (Kishida *et al*., 2022), or fluorescent MS2 (Harb *et al*., 2020) or R17 (Clarke *et al*., 2008) phages to label the F pilus, thus validating our labeling procedure and analysis methods.

Using the data of 647 pili detections originating from 387 donor cells allowed us to construct a 2D density map illustrating the localization of F pili around the cells (Fig. 1F). We observe that pili are anchored throughout the entire cell periphery, with a notable preference for the proximity of the midcell region, while anchoring at the cell tip are much less frequent (Fig. 1G). This positioning pattern remains consistent throughout the growth of the cells, as it is equally observed in both smaller (< 4 µm) and larger (≥ 4 µm) cells (Fig. S1E-1H). Pili detection also provides insights into their flexible nature *in vivo*. When the pili are shorter than 0.6 µm, they project orthogonally and straight from the cell surface. However, as the pili grow longer, they progressively exhibit a more pronounced apparent curvature (Fig. 1H). The number, length, and curvature of pili likely are key factors that determine the donor cell’s ability to probe its immediate surroundings. The 2D density map depicting the localization of pili enables us to delineate a three-dimensional (3D) space encompassing the donor cell that will be efficiently explored by the pili tips (dashed orange line in Fig. 1F). Analysis reveals that the majority of pili tips (594 out of 643) are confined within a 137.4 µm³ volume, facilitating highly efficient probing within this region. Outside of this volume, probing becomes dependent on the relatively infrequent occurrence of larger pili (53 out of 643), and is therefore predicted to be much less efficient.

### F pilus dynamics

Next, we performed time lapse imaging in a microfluidics chamber to visualize the real-time dynamics of F pili at the surface of donor cells in the absence of recipient cells. Using acquisition with 1 or 3 min/frame intervals, we observe that the number of pili per cell varies rapidly (Movie S1 and S2). Cells that are initially unpiliated on the first frame of the time lapse are able to produce pili within minutes (Movie S2), suggesting that the most if not all donor cells are able to produce pili. Some pili appear or disappear between two consecutive frames, while others remain extended for more than 60 min (Movie S3). To capture the dynamics of pilus extension and retraction with better time resolution at the single-cell level we acquired images with 10 sec/frame intervals (Fig. 2A; Movie S4). Analysis of isolated extension and retraction events reveals an average 49.5 ± 50 nm/s (n= 238) extension rate, remarkably similar to the average 47.5 ± 45 nm/s (n = 275) retraction rate (Fig. 2B; Movie S5). These results differ from those observed with F pili labelled with fluorescent R17 phages, which exhibited a retraction rate (15 nm/s) much slower than the extension rate (39 nm/s), likely accounting for the alteration of the pilus retraction by the binding of the phage itself (Clarke et al., 2008). Live-cell imaging also exposes the rapid switching between pili extension and retraction (Movie S4). Notably, repetitive extension-retraction events are frequently observed at specific positions along the cell membrane, suggesting that multiple pili can be successively produced by the same T4SS (Movie S4). This observation raised the question if reversal events occur at a specific pilus size or with a definite time periodicity. To address this question, we measured the length of the pili when they switch from extension to retraction (Fig. 2C). We observed that the distribution of pili lengths at the onset of retraction is not statistically different to that of the whole pili population (compare Fig. 2C to Fig. 1E), indicating that retraction occurs at any pilus length. Next, we measured an average 26 ± 13 seconds (n = 215) time lag between extension and retraction events, with widespread individual data points (Fig. 2D). These findings suggest that the events of pilus reversal exhibit significant variability in terms of both length and timing. This supports the notion that the transitions from extension to retraction during pilus biogenesis are more likely to be random rather than precisely regulated by an internal clock.

**Figure 2.**
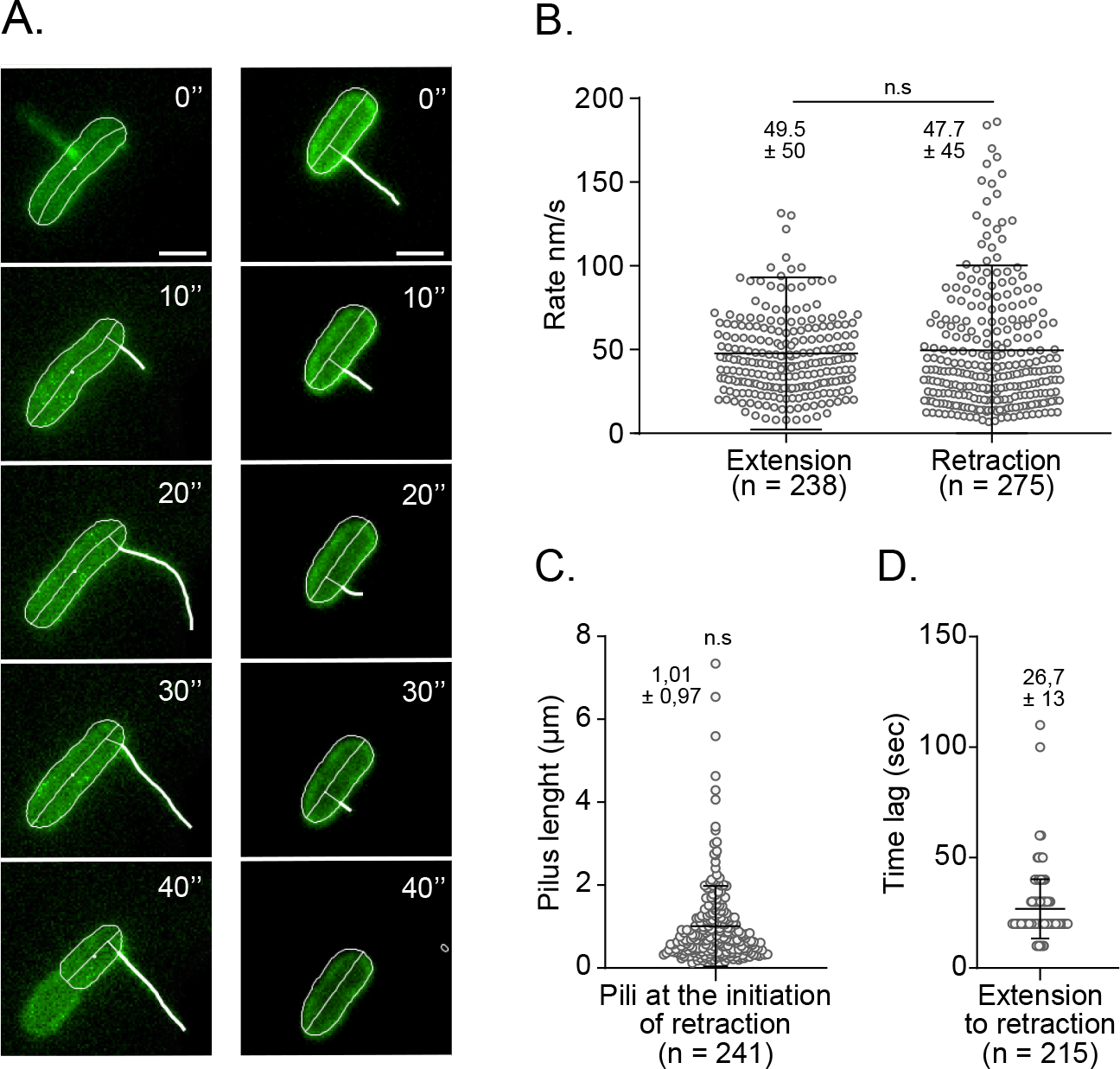
Extension and retraction dynamics of the F pilus (**A**) Time-laps microscopy image (10 sec/frame) of dynamic extension (on the left) and retraction (on the right) of the F pili. The contour made by the automatic detection of the pluging in MicrobeJ is represented in white. Scale bar 1 µm. (**B**) Jitter plot representing the speed of extension and retraction of the F pili in nm/s. Retraction and extension events analysed using 10 sec/frame time-laps. The Mean and SD was calculated from the indicated number of retraction/extension events. *P-value* significance from Mann-Whitney two-sided statistical test is indicated by n.s (non-significant, P = 0.13). (**C**) Distribution of the pili length at the initiation of a retraction. The Mean and SD was calculated from the indicated number of pili reversal events. *P-value* significance from Mann- Whitney two-sided statistical test compared the length distribution of all pili (Fig. 1E) is indicated by n.s (non-significant, P = 0.14). (**D**) Dot-plot showing the time lag (sec) between extension and retraction events. The Mean and SD was calculated from the indicated number of reversal events.

All together, these observations indicate that the heterogeneity observed in the number and length of pili within the donor population reflects a dynamic equilibrium resulting from the rapid processes of extension, retraction, and reversal events at the single-cell level. The highly dynamic nature of pilus biogenesis, coupled with the inherent flexibility of the pili, likely serves to optimize the probing ability of the donor cells in their search for a suitable attachment surface. It is worth noting that even in the absence of recipient cells, we consistently observe attachment of the pilus to the PDMS surface of the microfluidic chamber (Movie S6), as well as attachment to other donor cells (Movie S7) resulting in the formation of cells aggregates (Movie S8). We can note that attachment events are not necessarily followed by pilus retraction. These observations highlight the sticky property of the pili tips, enabling them to interact with biotic as well as abiotic surfaces.

### DNA transfer through extended F pili

To characterize the dynamics of the F pilus during conjugation, we combined F pilus labelling with reporters systems previously developed to monitor plasmid transfer using real-time microscopy (Nolivos *et al*., 2019; Goldlust *et al*., 2022; Couturier *et al*., 2023). The precise moment of ssDNA plasmid transfer from donor to recipient was captured using a fluorescent protein fusion of the chromosomally encoded single-strand-binding protein Ssb (Ssb-YPet). During conjugation, Ssb- YPet forms bright membrane-associated foci revealing the presence of single-strand plasmid DNA on both sides of the conjugation pore (Couturier *et al*., 2023). The conversion of the ssDNA plasmid into dsDNA is a hallmark for successful acquisition of the plasmid by the recipient cell and can be visualized by binding of mCherry-ParB (mCh-ParB) to a dsDNA *parS* site located on the plasmid resulting in a fluorescent focus formation (Nolivos *et al*., 2019; Goldlust *et al*., 2022; Couturier *et al*., 2023).

We performed time lapse microscopy imaging (3 min/frame intervals) of conjugation mixes between TraA^S54C^-producing *E. coli* F donors pre-labelled with maleimide (AF-Mal^488^ or AF-Mal^594^) and recipient cells (Fig. 3A-3C). Loading of the conjugation mix within the microfluidic chamber results in microscopy fields of view with a mono-layer of bacteria where most cells are in contact with other cells. Consequently, the vast majority of conjugation events observed occur between donor and recipient cells that are in tight cell-to-cell contact. The dynamics of the ssDNA transfer between cells in direct contact has been previously described by Couturier et al (Couturier *et al*., 2023). In the present study, we focused on conjugation events where donor and recipient cells were physically distant but connected via an extended F pilus.

**Figure 3.**
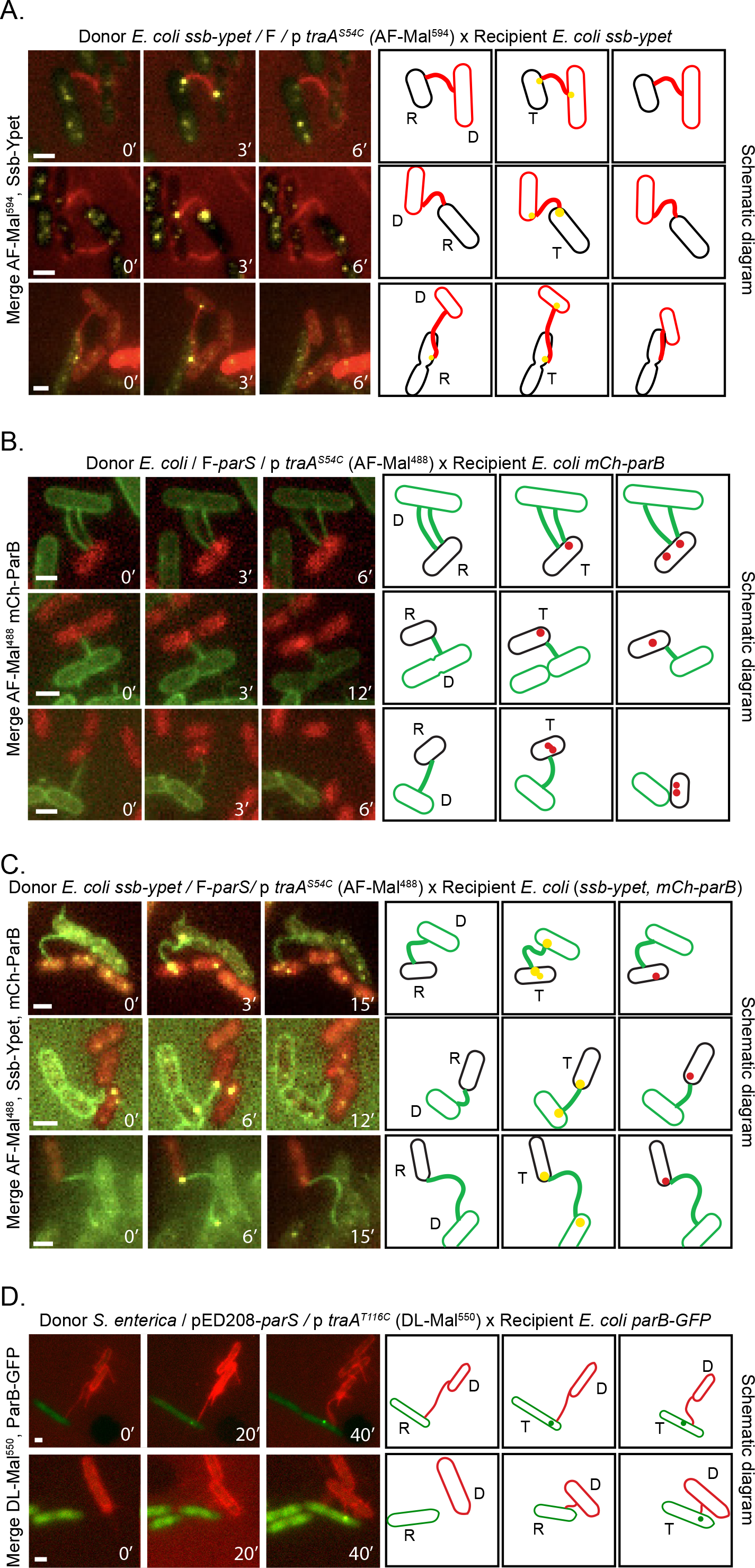
Plasmid transfer between physically distant cells (**A**) Time laps images of distant plasmid transfer events between *E. coli ssb-ypet* / F / p *traA*^S54C^ donors labelled with AF594-Mal and *E. coli ssb-ypet* recipient cells. Distant transfer is reported by the simultaneous formation of bright membrane-proximal Ssb-Ypet conjugative foci at each edge of the pilus connecting the mating pair cells. (**B**) Time-laps images of distant transfer events between *wt* A. *coli* / F*-parS* / p *traA*^S54C^ donors labelled with AF488-Mal and *E. coli* recipients producing mCh- ParB. Distant transfer is reported by the formation of mCh-ParB foci confirming the acquisition of the dsDNA plasmid by the recipient cell. (**C**) Time-laps images of distant transfer events between *wt E. coli ssb-ypet* / F*-parS* / p traA^S54C^ donors labelled with AF488-Mal and *E. coli ssb-ypet* recipients producing mCh-ParB. Distant transfer is reported by both the formation Ssb-Ypet conjugative foci at each edge of the pilus connecting the mating pair cells and the subsequent the formation of mCh- ParB foci, which confirms the acquisition of the dsDNA plasmid by the recipient cell. (**D**) Time-laps images of distant transfer events between *S. enterica / pED208 / p traA*^T116C^ donor cells labelled with DL550-Mal and *wt E. coli* recipient cells production ParB-GFP. (**A**), (**B**), (**C**) and (**D**) Scale bar 1 µm.

First, we examined Ssb-Ypet dynamics in recipients and donors labelled with AF-Mal^594^ to co-visualise the transferred ssDNA plasmid together with the F pilus (Fig. 3A, Fig. S2A; Movie S9). Analysis of the time-lapse images reveales distant ssDNA transfer events as determined by the simultaneous formation of bright membrane-associated Ssb-Ypet conjugative foci in the mating pair cell. These Ssb-Ypet foci are precisely located at both ends of the pilus, corresponding to the anchoring and the attachment point of the pilus on the donor’s and the recipient’s surface, respectively. The Ssb-Ypet foci were observed in one frame only, consistently with the 2.9 min lifespan of the ssDNA previously reported (Couturier *et al*., 2023). Next, we combined a F-*parS* donor labelled with AF-Mal^488^ and a recipient producing mCh-ParB to visualise the F pilus and the acquisition of the dsDNA plasmid by the recipient cell (Fig. 3B, Fig. S2B; Movie S10). Again, we observed that recipients, which were connected to a donor cell by an extended pilus, acquired the dsDNA plasmid as revealed by formation of a mCh-ParB focus. We note that mCh-ParB foci appeared in the vicinity of the pilus attachment point, consistent with the previous report that ssDNA- to-dsDNA conversion takes place at the location of plasmid entry (Couturier *et al*., 2023). Next, we performed three-color imaging using a *ssb-ypet* donor labelled with AF-Mal^488^ and a recipient strain producing both Ssb-Ypet and mCh-ParB (Fig. 3C, Fig. S3C; Movie S11). This experiment allowed us to visualise both the ssDNA transfer through the extended pilus, and the conversion of the ssDNA into dsDNA within the recipient, thus ascertaining a successful conjugation event through the pilus between cells that are not in tight contact but just bridged by an extended conjugative F pilus. Interestingly, we observed that long-distance plasmid transfer can occur through both straight and bent pili (Fig. 3).

Finally, we investigated whether the ability to transport DNA between physically distant cells was conserved for other F-like plasmids and during inter-species DNA transfer. Following the same experimental strategy than with *E. coli* F donors, we constructed a *Salmonella enterica* strain carrying the F-like plasmid pED208, a derepressed derivative plasmid of the F0Lac family (IncFV), first identified in *Salmonella* Typhi (Finlay *et al*., 1983; Lu *et al*., 2002), harboring a *parS* binding site and an expression plasmid (p *traA*^T116C^) coding for a mutant TraA^T116C^ pilin for bioconjugation with fluorescent Maleimide (Fig. S3A) (Kishida *et al*., 2022). This donor strain enabled efficient fluorescent labelling of pED208-encoded pili and exhibited *wt* conjugation efficiency a (Fig. S3B- 3C). Time-lapse imaging of pED208-*parS* conjugation in the microfluidic chamber was performed using a ParB-GFP-producing *E. coli* recipient strain (Fig. 3D). Analyses of ParB-GFP during time- lapse microscopy imaging revealed that *E. coli* recipient cells, which were only connected to an *S. enterica* donor cell by an extended pilus, acquired the dsDNA pED208 plasmid, This result supports that distant DNA transfer through the extended pilus is a conserved mechanism during conjugation of F-like plasmids (Fig. 3D).

## Discussion

In this study, we used maleimide bioconjugation to fluorescently label and investigate spatiotemporal dynamics of the F pilus in living cells, thus providing valuable insights into the DNA transfer mechanism during bacterial conjugation. F pili were found to be anchored unevenly on the side of the donor cells, primarily near the midcell position and rarely in the polar region. This localisation is remarkably reminiscent of the position of exit of the single-stranded DNA from donor cells during F plasmid transfer (Couturier *et al*., 2023), indicating that T4SS, which are active in pilus biogenesis and DNA transfer, are specifically rather than randomly positioned along the cell surface. This specific positioning pattern could reflect the non-random spatial arrangement of the T4SS machinery of the F plasmid within the cell, which is currently undocumented. Alternatively, if the T4SS machinery of the F plasmid were uniformly distributed along the cell periphery, similar to what has been observed for the pTi and R388 plasmids (Aguilar *et al*., 2011; Carranza *et al*., 2021), this pattern would strongly support the hypothesis that only the lateral T4SS pores are activated. This activation could be facilitated by the increased accessibility to F plasmid molecules, which are located at quarter positions within the cell and excluded from the cell poles (Niki and Hiraga, 1997; Gordon *et al*., 2004). Our results also corroborate previous studies using fluorescent phages (Clarke et al., 2008; Harb et al., 2020), demonstrating that the F pilus undergoes cycles of extension and retraction, even in the absence of recipient cells. Reversal events occur abruptly and randomly at any pilus length, with a highly variable time pattern. It is worth noting that the extension speed of the F pilus (49.5 nm/s) is relatively slow compared to other pili systems, such as *Vibrio cholerae* competence pilus (90 nm/s) (Ellison et al., 2018) or the Tad Pili of *Caulobacter crescentus* (140 nm/s) (Ellison et al., 2017). The moderate extension speed, combined with random extension-retraction cycles, pilus multiplicity, length, and flexibility, likely contribute to optimizing the ability of the F pilus to explore the cell’s environment and increasing the probability of contacting a recipient cell.

Importantly, our results show that pili encoded by the F and the pED208 F-like plasmids can serve as a conduit for DNA transport during conjugation between donor and recipient cells that are physically distant, thereby answering a long-standing and controversial question raised by Brinton almost 60 years ago (Brinton et al., 1964; Brinton, 1965). This novel observation was possible thanks to the development of specific fluorescent reporter systems but also to the use of the microfluidics apparatus that offers several advantages. Unlike agarose-mounted microscopy slides, the microfluidic setup immobilized bacterial cells while allowing the pili to move freely on the cell sides. In addition, it was first reported by Dürrenberger et al. (Dürrenberger *et al*., 1991) and later confirmed by Clarke et al. (Clarke et al., 2008) that the pilus retraction is associated with the rotation of the donor cell in relation to the recipient cell. However, cells immobilized within the microfluidic chamber are not able to rotate freely, which we think was critical to impede pilus retraction and enrich the microscopy fields of view with configurations where distant donor and recipient cells are connected solely by an extended pilus. Accordingly, this experimental setup provides the first direct observation of distant transfer via the F pilus, which was previously suggested by indirect evidence only (Harrington and Rogerson, 1990; Babić et al., 2008). Our findings also substantiate several inferences in favor of distant transfer made from structural data. The F pilus lumen diameter (28 Å) is large enough to accommodate for the passage of the single-stranded DNA molecule attached to the TraI relaxases, which is unfolded during the passage through the T4SS (Arutyunov and Frost, 2013; Trokter and Waksman, 2018). Furthermore, while the exact confirmation of the T4SS during transfer (Hu *et al*., 2019) and the route of the DNA through the T4SS machinery remains largely elusive, recent discoveries support the idea that the physico-chemical environment within the lumen is compatible with DNA transport. Indeed, work by Costa et al. (Costa *et al*., 2016) demonstrates that the alpha2- alpha3 loops of the TraA pilin project towards the lumen of the pilus, and would be in contact with the translocated DNA. They also demonstrate that each TraA pilin composing the F pilus is associated with a phosphatidylglycerol (PG) molecule, the head of which is exposed to the pilus limen. These PG molecules are proposed to render the lumen’s electrostatic potential electronegative, thereby facilitating the transport of the ssDNA through the pilus.

Our work allows us to propose a new model for bacterial conjugation. We suggest refining the term “contact-dependent” traditionally used to depict DNA conjugation, by introducing the terms “tight transfer” and “distant transfer”. Tight transfer occurs either when bacteria are initially in cell- to-cell contact or after pilus attachment to the recipient cell resulted in complete retraction. Tight transfer is likely the most efficient conjugation pathway, as the establishment of cell-to-cell contact is followed by the stabilization of shearing force-resistant mating pairs through the interaction between the TraN protein exposed on the donor cell surface and the recipient’s outer membrane protein OmpA (Low *et al*., 2022). Alternatively, when pilus attachment cannot be followed by complete retraction, distant transfer may occur through an extended F pilus. The balance between tight and distant transfer would likely be influenced by factors such as medium properties and cell density, both in laboratory conditions and in natural ecosystems. Enterobacterial carrying F-like plasmids primarily resides in the intestines of vertebrates, where they are surrounded by viscoelastic intestinal mucus that constrains bacterial movement. In these conditions, distant transfer through extended pili is likely physiologically relevant to compensate for the inability to establish direct contact between the mating cells. Whether the ability to transport DNA is restricted to specialised pili, such as thin and flexible pili encoded by F-like plasmids, or whether this is a general property of conjugative pili remains to be addressed.

The existence of both tight and distant transfer has implications regarding the regulation and molecular mechanism of F-like plasmids conjugation. It is well established that DNA transfer is triggered upon contact with the recipient bacterium, via a signal which nature remains unknown. While our findings do not uncover the exact nature of this signal, they do establish that wall-to-wall contact between donor and recipient cells is not essential for initiating DNA transfer. Instead, it is conceivable that the signal for transfer initiation is transmitted through a pilus-dependent mechanosensing mechanism, activated either by complete pilus retraction or when pilus retraction is hindered. Additionally, although our data demonstrate the ability of pili to transport DNA, existing literature consistently reports that extended F pili are dispensable for conjugation when stable wall-to-wall cell interactions are present. Donors lacking detectable pili can still perform plasmid transfer and often exhibit efficient conjugation under solid conditions (Panicker and Minkley, 1985) (Kishida et al., 2022). This indicates that the production of surface-exposed pili is not required for transfer, as long as stable donor-recipient contact is facilitated. While extended pili are not required for conjugation, the TraA pilin subunit remains essential for conjugation in all tested conditions. The existence of TraA fibers that are not extended extracellularly but have exposed tips at the cell surface has been proposed from early experiments by Novotny and Fives-Taylor (Novotny and Fives-Taylor, 1974). They demonstrated that pili disassembly induced by NaCN hinders the adsorption of R17 bacteriophage to the sides of the pili but do not affect the adsorption of M13 phage to the pili tips. Therefore, it is possible that, even during tight transfer between closely apposed cells, the presence of very short “vestigial” pili is necessary for conjugation. Such rudimentary TraA polymers located within the T4SS system spanning the periplasm could make contact with the recipient cell’s surface and potentially play a critical role in transporting the DNA through the conjugation pore during transfer.

## Acknowledgements

The authors thank the National BioResource Project and Coli Genetic Stock Center for providing strains, Tiago R.D. Costa for providing pED208 and the *wt* p *traA* plasmid and Nelly Schropp for early involvement in the project and help in plasmid construction.

## Author contributions

K.G. and M.H. performed the experiment analysed the data and prepared the figures; A.D. developed the MicrobeJ plugin for pili automatic detection; A.B-D. constructed strains; M.E. provided supervision, revised the paper, and provided funding, C.L. conceived and designed the project, wrote the paper, edited the figures, and provided funding.

## Data and materials availability

All data to understand and assess the conclusions of this research are available in the main text and Supplementary Materials.

## Methods

### Bacterial strains, plasmids, and growth culture conditions

Bacterial strains are listed in Table S1, plasmids in Table S2, and oligonucleotides in Tables S3. Construction of the bacterial strains and plasmid for the F-pilus labelling are described in Goldlust et al. (Goldlust *et al*., 2022). Briefly, to constructs TraA single-mutant proteins (TraA^S54C^) of the serine S54 substituted by a cysteine, first traA gene was inserted into the pTrc99a expression plasmid by Gibson assembly and verified by Sanger sequencing (Eurofins Genomics biotech). Then, cysteine substitution by in vivo assembly mutagenesis was performed directly on the pTrc99a *traA*. Fusion of genes with fluorescent tags used λRed recombination (Yu *et al*., 2000; Datsenko and Wanner, 2000). F plasmids were transferred by conjugation into a background strain K12 MG1655. When necessary the *kan* and *cat* genes were removed using site-specific recombination induced by expression of the Flp recombinase from plasmid pCP20 (Datsenko and Wanner, 2000). Plasmid cloning were done by Gibson Assembly and verified by Sanger sequencing (Eurofins Genomics biotech). Cells were grown at 37°C in Luria-Bertani (LB) broth medium and wash in Rich defined medium (RDM) before imaging. When appropriate, supplements were used at the following concentrations; Streptomycin (St) 20 µg/ml, Ampicillin (Amp) 100 µg/ml, Kanamycin (Kn) 50 µg/ml and Tetracycline (Tc) 10 µg/ml and IPTG 40 µM.

### Conjugation assays

Overnight cultures in LB of recipient and donor cells were diluted to an A_600_ of 0.05 and grown until an A_600_ comprised between 0.7 and 0.9 was reached. 25 µl of donor and 75 µl of recipient cultures were mixed into an Eppendorf tube and incubated for 60 minutes at 37°C. Conjugation mix were then vortexed, serial diluted, and plated on LB agar supplemented the appropriate antibiotic to select for donor, recipient or transconjugant populations.

### Live-cell microscopy experiments and image acquisition

For all experiments, overnight cultures in LB were diluted to an A600 of 0.05 and grown until A600 = 0.4. Donor cells are incubated 30 min with Alexa Fluor 488 C5-maleimide (AF-Mal^488^) or AF594 C5-maleimide (AF-Mal^594^) then wash in RDM before microscopy observation.

*Time lapse experiments into microfluidics chambers.* Time lapse imaging in a microfluidic chamber is performed as previously described (Nolivos *et al*., 2019; Couturier *et al*., 2023; Cayron *et al*., 2023). Conjugation sample were obtained by mixing 50 µl of the pre-labelled donor and 50 µl of recipient into and Eppendorf tube, then 50 µl of conjugation mix was loaded into a B04A microfluidic chamber (ONIX, CellASIC®). The temperature was maintained at 37°C during imaging. For donor alone, cells were imaged every ten seconds, one or three minutes. For conjugation imaging, conjugation mixes were imaged every 3 or 20 minutes.

*Snapshot microscopy in agarose pads for characterisation of donor alone.* Agarose-mounted slides are prepared as in Lesterlin and Duabrry, 2016. 10 µl samples of clonal culture were spotted onto an RDM 1% agarose pad on and imaged directly into thermostatic chambers at 37°C.

*Image acquisition.* Conventional wide-field fluorescence microscopy imaging was carried out on an Eclipse Ti2-E microscope (Nikon), equipped with x100/1.45 oil Plan Apo Lambda phase objective, ORCA-Fusion digital CMOS camera (Hamamatsu), and using NIS software for image acquisition. Acquisitions were performed using 50% power of a Fluo LED Spectra X light source at 488 nm and 560 nm excitation wavelengths. For time-laps experiment, exposure settings were 100 ms for Ypet, AF594-Mal, AF488-Mal, and mCherry, 50 ms for phase contrast. For snapshots experiments exposure settings were 300 ms for AF594-Mal and AF488-Mal, 100 ms for Ypet, and mCherry.

### Image analysis

Quantitative image analysis was done using Fiji software with MicrobeJ plugin (Ducret et al., 2016). For snapshot analysis on donor allowing the characterisation of the F pili, cell’s outline and pili detection was performed automatically using MicrobeJ custom plugin. Error of detection was corrected manually using the editing interface. Pili localisation parameters were automatically extracted and plotted using MicrobeJ. For time lapse experiments, detection of cells was done semi- automatedly using the Manual-editing interface and pili detection was performed automatically using MicrobeJ custom plugin. For characterisation of donor alone, pili length (µm) and number per cell were extracted automatically. Plots presenting time lapse data for dynamics of F pili alone were either aligned to the first frame where the cell exhibits a F pili retraction or extension or as indicated in the corresponding figure legend.

### Statistical analysis

P-value significance were analysed running specific statistical tests on the GraphPad Prism software. Pilus length, extension and retraction data from quantitative microscopy analysis were extracted from the MicrobeJ interface and transferred to GraphPad. P-value significance was then performed using unpaired non-parametric Mann-Whitney two-sided statistical test. When required, P-value and significance are indicated on the figure panels and within the corresponding legend.

### Data availability

All data to understand and assess the conclusions of this research are available in the main text and Supplementary Information. Source data are provided with this paper in the Source- Data file. Raw microscopy data will be available on Figshare at the time of publication.

