## Supplementary information for "The F pilus serves as a conduit for the DNA during conjugation between physically distant bacteria"

**This file includes:**

Figures S1 to S3
Tables S1 to S3
Supplementary movie legends

**Other Supplementary Information for this manuscript include the following:**
Movies S1 to S11

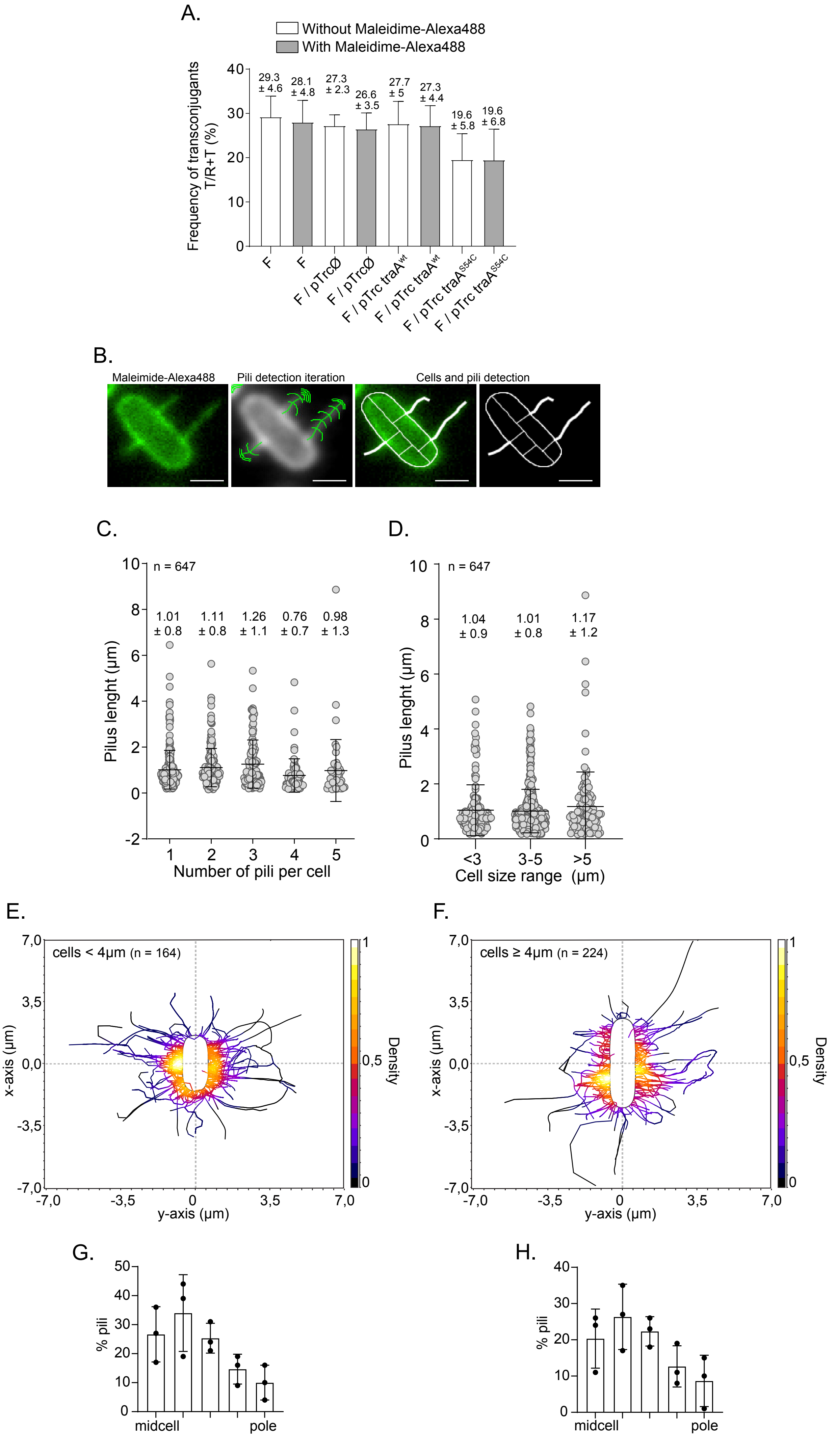

A.
Without Maleidime-Alexa488
With Maleidime-Alexa488
Frequency of transconjugants T/R+T (%)
29.3 ± 4.6
28.1 ± 4.8
27.3 ± 2.3
26.6 ± 3.5
27.7 ± 5
27.3 ± 4.4
19.6 ± 5.8
19.6 ± 6.8
F
F
F / pTrcØ
F / pTrcØ
F / pTrc traA^wt
F / pTrc traA^wt
F / pTrc traA^S54C
F / pTrc traA^S54C
B.
Maleimide-Alexa488
Pili detection iteration
Cells and pili detection
C.
n = 647
1.01 ± 0.8
1.11 ± 0.8
1.26 ± 1.1
0.76 ± 0.7
0.98 ± 1.3
Pilus lenght (µm)
Number of pili per cell
D.
n = 647
1.04 ± 0.9
1.01 ± 0.8
1.17 ± 1.2
Pilus lenght (µm)
<3
3-5
>5
Cell size range (µm)
E.
cells < 4µm (n = 164)
x-axis (µm)
y-axis (µm)
Density
F.
cells ≥ 4µm (n = 224)
x-axis (µm)
y-axis (µm)
Density
G.
% pili
midcell
pole
H.
% pili
midcell
pole


Figure S1

**Figure S1.**

(**A**) Histograms showing the conjugation efficiency of different *wt E. coli* / F donor strains. The Mean and SD was calculated from three independent biological replicates. (**B**) Microscopy image showing the process of automatic pili detection by MicrobeJ. The first image shows the AF488-Mal fluorescence channel. The second shows the process of pili detection by identification of the pilus anchoring point followed by iteration of pilus filament detection. The third and fourth image show the resulting detection of the cell outline and the pili in white. Scale bar 1 µm. (**C**) Jitter plot of the correlation between the pili length and the number of pili per cells. Each dot represent an individual pilus. The Mean and SD was calculated from the indicated number of pili. (**D**) Jitter plot of the correlation between the pili length and the cell size range. Each dot represent an individual pilus. The Mean and SD was calculated from the indicated number of pili. (**E**) & (**F**) 2D density map of pili localisation around the donor cell categorized be cell size (< and ≥ to 4 µm). The number of pili (n) from three biological replicates is indicated. Density scale on the left. (**G**) and (**H**) Histogram of pili distribution along the cell perimeter from midcell to the cell tip of donor cells categorized be cell size (< and ≥ to 4 µm).

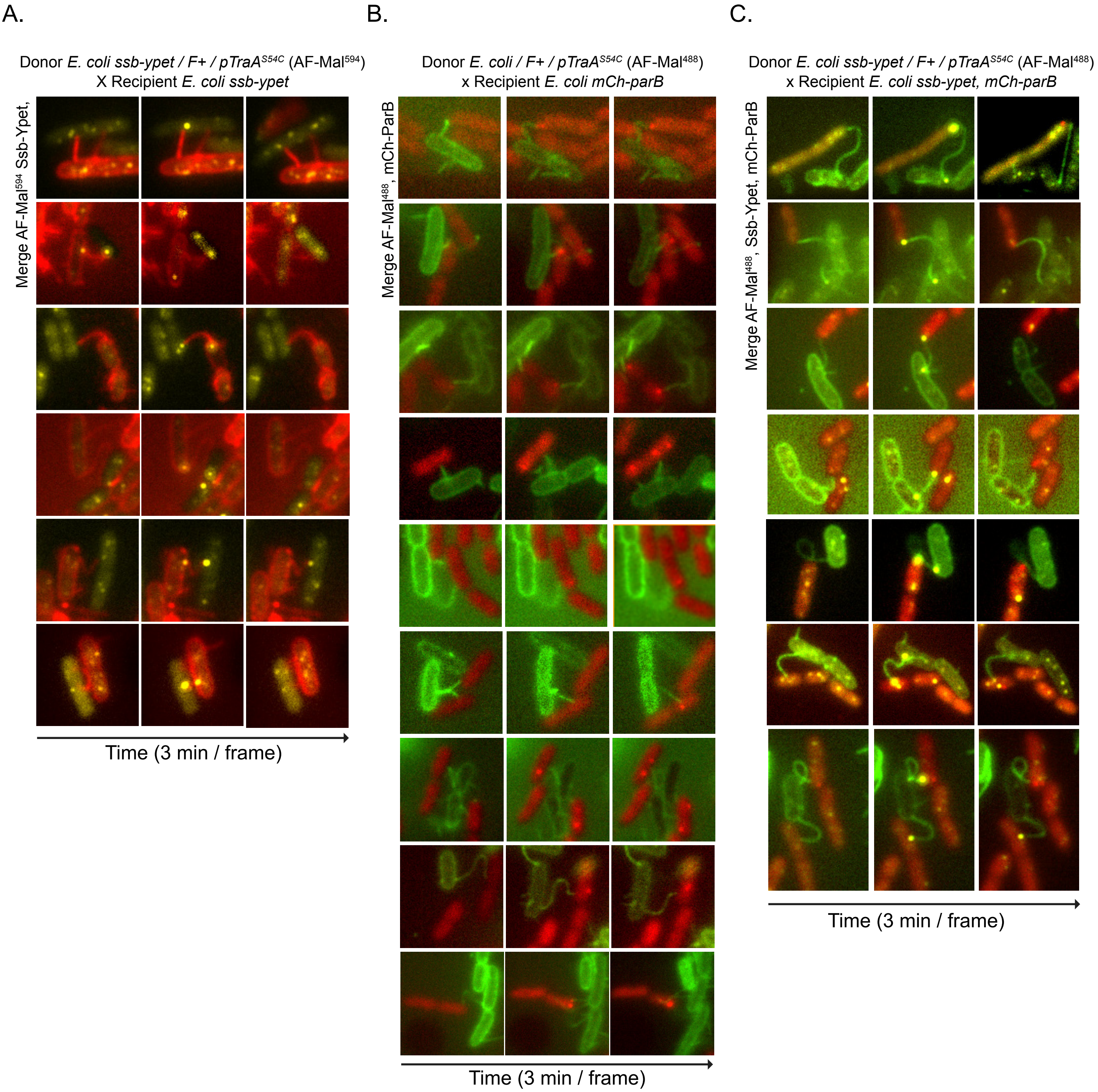

A.
Donor E. coli ssb-ypet / F+ / pTraA^S54C (AF-Mal^594)
X Recipient E. coli ssb-ypet
Merge AF-Mal^594 Ssb-Ypet,
Time (3 min / frame)
B.
Donor E. coli / F+ / pTraA^S54C (AF-Mal^488)
x Recipient E. coli mCh-parB
Merge AF-Mal^488, mCh-ParB
Time (3 min / frame)
C.
Donor E. coli ssb-ypet / F+ / pTraA^S54C (AF-Mal^488)
x Recipient E. coli ssb-ypet, mCh-parB
Merge AF-Mal^488, Ssb-Ypet, mCh-ParB
Time (3 min / frame)


Figure S2

**Figure S2. Plasmid transfer between physically distant cells**

(**A**) Stills images from time lapses acquisition of distant transfer events between *E. coli ssb-ypet* / F / p *traA*$^{S54C}$ donors labelled with AF594-Mal and *E. coli ssb-ypet* recipient cells. Distant transfer is reported by the simultaneous formation of bright membrane-proximal Ssb-Ypet conjugative foci at each edge of the pilus connecting the mating pair cells. (**B**) Stills images from time lapses acquisition of distant transfer events between *wt E. coli* / F-*parS* / p *traA*$^{S54C}$ donors labelled with AF488-Mal and *E. coli* recipients producing mCh-ParB. Distant transfer is reported by the formation of mCh-ParB foci confirming the acquisition of the dsDNA plasmid by the recipient cell. (**C**) Stills images from time lapses acquisition of distant transfer events between *wt E. coli ssb-ypet* / F-*parS* / p traA$^{S54C}$ donors labelled with AF488-Mal and *E. coli ssb-ypet* recipients producing mCh-ParB. Distant transfer is reported by both the formation Ssb-Ypet conjugative foci at each edge of the pilus connecting the mating pair cells and the subsequent the formation of mCh-ParB foci, which confirms the acquisition of the dsDNA plasmid by the recipient cell. (**A**), (**B**) and (**C**) the time-laps acquired using 3 min/frame time intervals. Scale bar 1 µm.

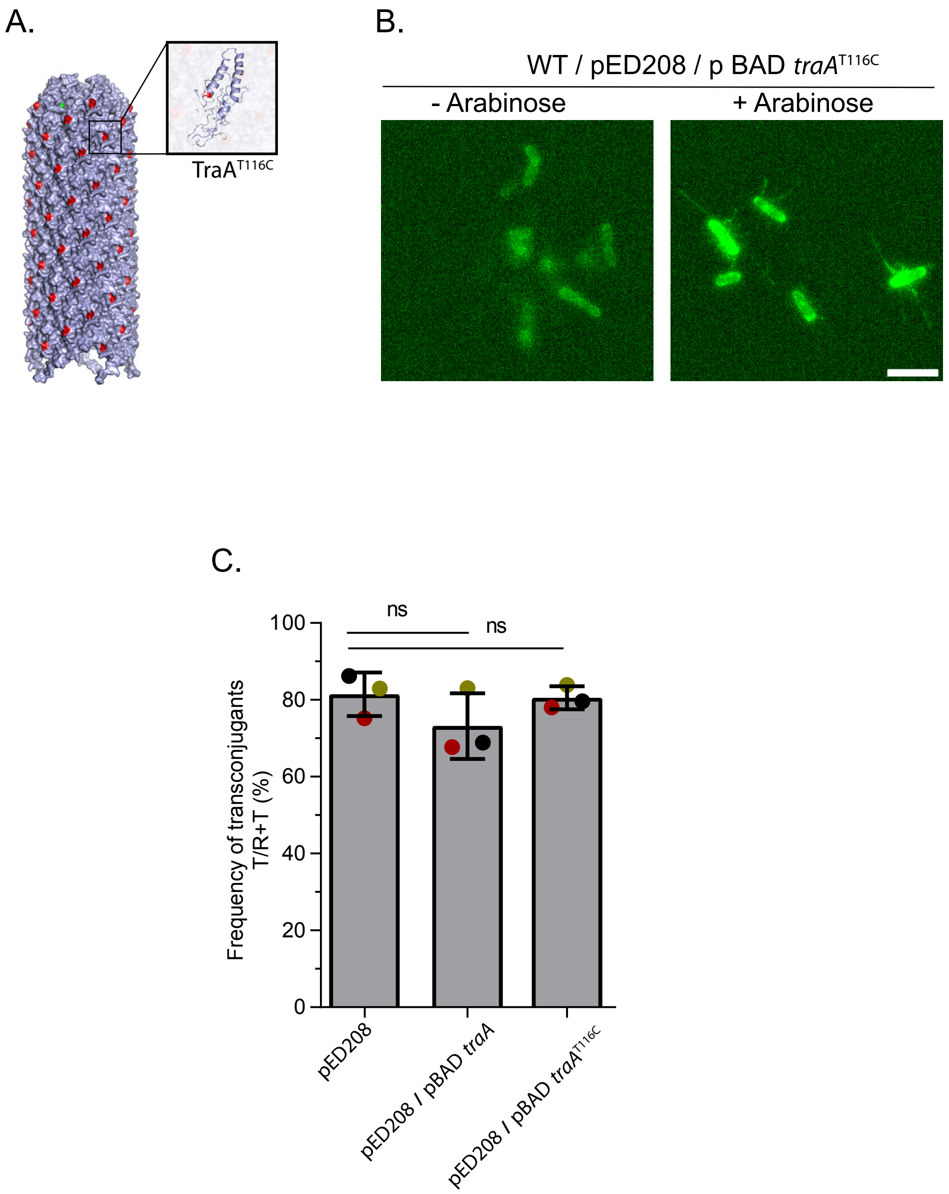
A.
TraA$^{T116C}$
B.
WT / pED208 / p BAD *traA*$^{T116C}$
- Arabinose
+ Arabinose
C.
ns
ns
100
80
60
40
20
0
Frequency of transconjugants T/R+T (%)
pED208
pED208 / pBAD *traA*
pED208 / pBAD *traA*$^{T116C}$

Figure S3

**Figure S3. Plasmid transfer between physically distant cells**

(**A**) Surface representation map of the cryo-EM structure of the pED208 conjugative pilus (adapted form (Costa *et al.*, 2016)) showing the surface-exposed Threonine 116 (T61 in the mature pilin) replaced by a cysteine in the TraA$^{T116C}$ mutant. The inset shows the structure of an individual TraA pilin with the T116C residue in red. (**B**) Microscopy image of *wt Salmonella enterica* cells carrying the pED208 and the pBAD *traA*$^{T116C}$ plasmids labelled with maleimide STAR Green. No labelled pili were observed in the absence of arabinose induction of TraA$^{T116C}$. Scale bar 1 µm. (**C**) Conjugation efficiency between different *S. enterica* / pED208 donors carrying or not the pBAD *traA* and pBAD *traA*$^{T116C}$ plasmids, and *wt E. coli* recipients, in the absence or the presence of the pBAD *traA* and pBAD *traA*$^{T116C}$ plasmids. The Mean and SD was calculated from three independent biological replicates.

| Strain | Relevant genotype[a, b, c] | Source or reference[a] |
|---|---|---|
| **MG1655 and derivatives** | | |
| **LY110** | MS388 / F-Tn*10, parS*$_{PMT1}$*-FRT-cat-FRT* | [1] |
| **LY117** | MS388 *ssb-ypet-FRT-kan-FRT* | [1] |
| **LY318** | MS388 / pSN70 | [1] |
| **LY358** | MS388 *ssb-ypet-FRT* / pSN70 | [1] |
| **LY2095** | MS388/ F-Tn*10, parS*$_{PMT1}$*-FRT-cat-FRT ssb-ypet-FRT-kan-FRT* | Conjugation LY110 x LY117 to Kn$^R$ Tc$^r$ |
| **LY1149** | MS388 / F-Tn*10, parS*$_{PMT1}$*-FRT-traC-sfGFP-FRT* | |
| **LY2103** | MS388 / F-Tn*10, parS*$_{PMT1}$*-FRT-cat-FRT / pTrc99a* | Transformation *pTrc99a* in LY110 |
| **LY2104** | MS388 / F-Tn*10, parS*$_{PMT1}$*-FRT-cat-FRT / pTrc99a-traA* | Transformation *pTrc99a-traA* in LY110 |
| **LY1500** | MS388 / F-Tn*10, parS*$_{PMT1}$*-FRT-cat-FRT / pTrc99a-traA-S54* | Transformation *pTrc99a-traA-S54C* in LY110 |
| **LY2099** | MS388/ F-Tn*10, parS*$_{PMT1}$*-FRT-cat-FRT/ ssb-ypet-FRT-kan-FRT / pTrc99a-traA-S54* | Transformation *pTrc99a-traA-S54C* in LY2095 |
| **LY2193** | MS388 / F-Tn*10, parS*$_{PMT1}$*-FRT-traC-sfGFP-FRT / pTrc99a-traA-S54* | Transformation *pTrc99a-traA-S54C in* LY1149 |
| **MS388** | MG1655 *rpsL* (St$^R$) | Gift from F. Cornet |
| **MS428** | MG1655 *rpsL* (St$^R$), ΔlacZ | Gift from F. Cornet |

**Table S1. Strains list**

| Name | Construct and Usage [a] | Source or reference |
|---|---|---|
| **p mCherry-*ParB* (pSN70)** | IPTG inducible expression of N-terminal fusion mCherry-$ParB_{PMT1}$ | 1 |
| **pTrc99a** | | |
| **pGOLD** | *pTrc99a-traA* | |
| **pGOLD3** | *pTrc99a-traA-S54C* | |
| **F-Tn*10* conjugative plasmid (from K603) derivatives** | | |
| **F*wt*** | F-Tn*10* with *$parS_{PMT1}$* inserted at the intergenic *ygeB-ygfA locus* | 1 |
| **F *traC-sfgfp*** | F-Tn*10 $parS_{PMT1}$* with *traC-sfgfp* translational fusion at the endogenous locus | |

**Table S2. Plasmids used in this study**

| Name | Sequence | Construct |
|---|---|---|
| **OL602** | GATCCTCTAGAGTCGACCTGCAGGC | *pTrc99a* amplification |
| **OL675** | CATGGTCTGTTTCCTGTGTG | |
| **OL694** | CACACAGGAAACAGACCATGAATGCTGT TTTAAGTGT | TraA amplification on F with flancking region for insertion in *pTrc99a* |
| **OL695** | GGTCGACTCTAGAGGATCTCAGAGGCCA ACGACGGCCA | |
| **OL696** | GCGATGGCCGCCGGCTGCAGTGGTCAGG ACCTGATGGCAAGC | IVA assembly for replacement of Serine 54 by cysteine |
| **OL697** | AGCCGGCGGCCATCGCCAGCTG | |
| **OL217** | CCGACATCATAACGGTTCT | PCR verification of pTrc99a-TraA-S54C constructed |
| **OL219** | GGCTGAAAATCTTCTCTCAT | |

**Table S3. PCR primers used for strains and plasmids constructions**

**Supplementary movie legends**

**Movie S1.**

Microfluidic time-lapse imaging showing *wt E. coli* / F / p traA$^{S54C}$ labelled with AF488-Mal (left) with the corresponding cell outlines and pili detections performed using MicrobeJ custom plugin (right). Scale bar 1 µm and time in minutes are indicated (1 min/frame).

**Movie S2.**

Microfluidic time-lapse imaging of *wt E. coli* / F / p traA$^{S54C}$ labelled with AF488-Mal showing a rapidly varying number of pili per cell (left). The merge of the fluorescence images and the corresponding cell outlines and pili detections is shown (left). Scale bar 1 µm and time in minutes are indicated (3 min/frame).

**Movie S3.**

Microfluidic time-lapse imaging of an *E. coli* / F / p traA$^{S54C}$ cell labelled with AF488-Mal that exhibit long-lasting pilus. Scale bar 1 µm and time in minutes are indicated (3 min/frame).

**Movie S4.**

Combination of five time-lapses of *E. coli* / F / p *traA*$^{S54C}$ cells labelled with AF488-Mal illustrating the heterogeneity in pili number per cell, length, extension and retraction dynamics, and life span. Scale bar 1 µm and time in minutes are indicated (10 sec/frame).

**Movie S5.**

Microfluidic time-lapses of *wt E. coli* / F / p *traA*$^{S54C}$ labelled with AF488-Mal showing one extension (top) and one retraction (bottom) event. The merge of the fluorescence images and the corresponding cell outlines and pili detections are shown (left). Scale bar 1 µm and time in minutes are indicated (10 sec/frame).

**Movie S6.**

Time-lapse imaging showing attachment of the donor cell pili to the PDMS surface of the microfluidic chamber. A merge of phase contrast and AF488-Mal fluorescence images is shown. Scale bar 1 µm and time in minutes are indicated (3 min/frame).

**Movie S7.**

Time-lapses showing two events of attachment of the donor cell pili to other donor cells within the microfluidic chamber. Scale bar 1 μm and time in minutes are indicated (3 min/frame). Phase contrast image (left) and AF488-Mal fluorescence image (right) are shown.

**Movie S8.**

Time-lapses showing the attachment of multiple pili between several donor cells that result in the formation of cell aggregates within the microfluidic chamber. Scale bar 1 μm and time in minutes are indicated (3 min/frame).

**Movie S9.**

Time-lapses of distant transfer events between *E. coli ssb-ypet* / F / p *traA*$^{S54C}$ donors labelled with AF594-Mal and *E. coli ssb-ypet* recipient cells. Distant transfer is reported by the simultaneous formation of bright membrane-proximal Ssb-Ypet conjugative foci at each edge of the pilus connecting the mating pair cells. Five different distant transfer events are shown. Scale bar 1 μm and time in minutes are indicated (3 min/frame).

**Movie S10.**

Time-lapses of distant transfer events between *wt E. coli* / F-*parS* / p *traA*$^{S54C}$ donors labelled with AF488-Mal and *E. coli* recipients producing mCh-ParB. Distant transfer is reported by the formation of mCh-ParB foci confirming the acquisition of the dsDNA plasmid by the recipient cell. Six different distant transfer events are shown. Scale bar 1 μm and time in minutes are indicated (3 min/frame).

**Movie S11.**

Time-lapses of distant transfer events between *wt E. coli ssb-ypet* / F-*parS* / p traA$^{S54C}$ donors labelled with AF488-Mal and *E. coli ssb-ypet* recipients producing mCh-ParB. Distant transfer is reported by both the formation Ssb-Ypet conjugative foci at each edge of the pilus connecting the mating pair cells and the subsequent the formation of mCh-ParB foci, which confirms the acquisition of the dsDNA plasmid by the recipient cell. Scale bar 1 μm and time in minutes are indicated (3 min/frame).
